# Variation analysis of fruit phenotypic traits of different provenances of ancient *Cinnamomum camphora* in Jiangxi Province of China

**DOI:** 10.1101/2025.02.12.637828

**Authors:** Shengying Wan, Jiao Zhao, Jie Ma, Jie Zhang, Qingqing Liu, Changlong Xiao, Jiexi Hou, Zhinong Jin

**Affiliations:** Nanchang Institute of Technology, Nanchang, Jiangxi, China; Jiangxi Provincial Engineering Research Center of Seed-Breeding and Utilization of Camphor Trees, Nanchang, Jiangxi, China

## Abstract

Exploring fruit phenotypic diversity of different provenances of ancient *Cinnamomum camphora* in Jiangxi Province of China, it will provide a theoretical basis for the collection, selection and improvement of germplasm resources. Taking the naturally distributed ancient *Cinnamomum camphora* in 45 counties of 11 cities in Jiangxi Province as the research object, nested ANOVA, Pearson correlation analysis, principal component analysis and other methods were used to statistically analyze 10 phenotypic traits such as fruit thousand-grain weight, fruit volume, fruit side diameter, fruit horizontal diameter, fruit vertical diameter, and peel thickness in different provenances. The resuit was as follows. (1) The fruit phenotypic traits of ancient *Cinnamomum camphora* in Jiangxi Province were rich in variation among and within provenances, and the variation among provenances was the main source of phenotypic variation. The mean value of phenotypic differentiation coefficient among provenances was 80.1%. The coefficients of variation of fruit phenotypic traits among different provenances was ranged from 2.64% to 25.45%, with the largest coefficient of variation for peel thickness. The mean value of coefficient of variation among provenances(9.65%) was larger than that within provenances(8.52%). (2) Significant or highly significant correlations were existed among the fruit phenotypic traits of ancient *Cinnamomum camphora* in Jiangxi Province. Fruit thousand-grain weight was significantly positively correlated with fruit volume, fruit side diameter, fruit horizontal diameter, fruit vertical diameter, peel thickness, fruit side diameter-fruit horizontal diameter ratio, fruit side diameter-fruit vertical diameter ratio, fruit horizontal diameter-fruit vertical diameter ratio (*P*<0.05), while fruit horizontal diameter and fruit vertical diameter were significantly negatively correlated with fruit thousand-grain weight to fruit volume ratio(*P*<0.05). (3) There was a significant correlation between fruit phenotypic traits of ancient *Cinnamomum camphora* and latitude, longitude and climate factors in Jiangxi Province. Among them, fruit thousand-grain weight and fruit volume were highly significant negatively correlated (*P*<0.01) with annual average sunshine duration and mean temperature in July, and highly significant positively correlated with mean temperature in January and mean altitude factor (*P*<0.01). (4) The cumulative variance contribution of the first three principal components in the principal component analysis reached 78.78%, and the fruit side diameter, fruit side diameter-fruit vertical diameter ratio, and fruit side diameter-fruit horizontal diameter ratio could comprehensively reflect the information and ranking of the 10 traits. (5) In the cluster analysis, it was divided into two groups A and B at the distance coefficient of 1.2, and further divided into different subgroups at the distance coefficient of 0.9. The fruit morphology indexes from subgroup B2 were bigger. In conclusion, fruit phenotypic traits of ancient *Cinnamomum camphora* in Jiangxi Province were rich in variation, and the variation of among provenances was more than within provenances. At the same time, there was a certain geographic variation pattern of the phenotypic traits of *Cinnamomum camphora*. The results would provide an important direction for the selection and breeding of excellent germplasm resources in Jiangxi Province, China.

## Introduction

*Cinnamomum camphora* belongs to the camphor family camphor genus, evergreen trees, mainly distributed in the southern region of China. It has an important ecological and economic utilization value with timber, landscaping, medicine and biochemical raw materials and other functions[1].Therefore, the study of *Cinnamomum camphora* germplasm resources will provide a theoretical basis for the selection and breeding of excellent materials.

Plant phenotype is the combination of various morphological features of a species, which is the result of the co-regulation of genes and environment[2–3]. Genetic diversity is the synthesis of genetic variation among provenances and within provenances[4], and the studies of phenotypic diversity are mainly focused on the pattern of phenotypic variation and distribution under different habitat conditions, which is an important part of genetic diversity[5]. After long-term natural selection and environmental pressure, the growth and development level of plants often undergoes irreversible changes, which in turn causes the variation of morphological features to produce new phenotypes[6], and the research on phenotypic trait diversity can be used to understand the mechanism of plant response to the ecological environment and the law of genetic variation. Numerous studies have shown that the phenotypic traits in forest trees such as *Idesia polycarpa* Maxim., *Elaeagnus mollis* Diels, *Acer* ginnala, *Canarium album* L. and *Melia azedarach* are different from each other in terms of their phenotypic traits[7–10]. The phenotypic traits of forest trees are rich in variation among different provenances, which provides a basis for selection of good species and rational zoning of fruits. Differences in phenotypic traits of forest trees are related to many factors such as latitude, longitude and climatic factors[11].

Wang et al.[12] found that longitude, mean altitude and frost-free period were the main reasons affecting the variation of phenotypic traits in the fruit of *Rhus chinensis* Mill., whereas the phenotypic variation of *Dendrocalamus latiflorus* Munro. was affected by longitude, latitude, annual mean temperature, and mean annual precipitation, etc.[13]. At present, the phenotypic studies on ancient *Cinnamomum camphora* mainly focus on the leaves[14], only limited in Sichuan, Zhejiang in China. Jiangxi Province is an important distribution area of ancient *Cinnamomum camphora*, and the fruit phenotypic variation pattern in this area is still unclear. Therefore, ancient *Cinnamomum camphora* in Jiangxi Province of China was taken to research, and nested ANOVA, Pearson correlation analysis, regression analysis was carried out to explore the diversity of fruit phenotypes as well as the geographic variation law, so as to provide references for breeding selection and germplasm resource conservation.

## Materials and Methods

### Overview of the collection site

In this study, 170 healthy growth plants with age more than 100 years were selected in December 2018 from 45 counties of 11 cities in Jiangxi province, China. The geographic location information and meteorological data of various sources were shown in Table 1.

**Table 1.**
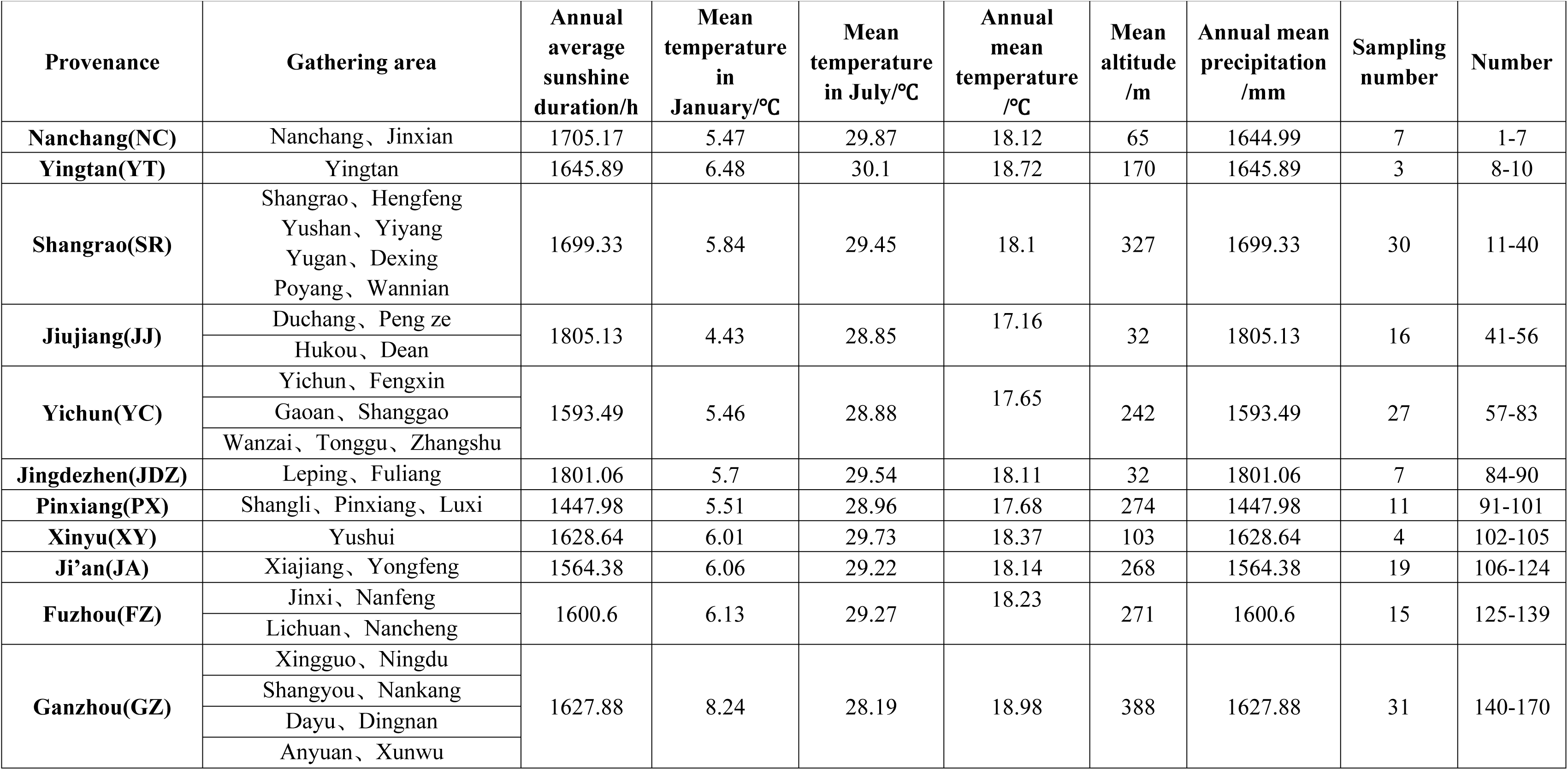
Geographic location information and sampling of ancient *Cinnamomum camphora* in different provenance.

### Data collection

Among the collected ancient *Cinnamomum camphora* fruits, 100 fruits were randomly selected from each tree, and the weight of each fruit was determined by electronic balance (precision 0. 0001g), and the volume of 100 grains of each fruit was measured by liquid difference method to calculate the fruit thousand-grain weight(FTW, g) and fruit volume(FV, cm^3^). 30 from the 100 fruits were selected to measure the fruit vertical diameter(FVD), fruit horizontal diameter(FHD), the fruit vertical diameter(FVD) and peel thickness(PT) of each fruit with electronic vernier caliper (precision 0.01mm). FVD was the maximum distance from the top to the base of the respective longitudinal section. FHD was the maximum distance from the ventral suture of the respective cross section to the other side, and FSD was the maximum distance on both sides of the respective ventral suture. Then, FSD/FHD, FSD/FVD, FHD/FVD and FTW/FV were calculated according to the measurement results.

### Data and statistics

Nested ANOVA, Pearson correlation analysis, regression analysis, and cluster analysis were performed using SPSS 26.0 software.

The coefficient of variation formula was given as follows, 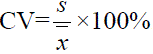, Where CV was the coefficient of variation, and xis the trait mean, S is the trait standard deviation.

Shannon-Wiener is a quantitative representation of the variation of trait diversity. The calculation formula was 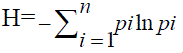. In the formula, H was the diversity index, and *Pi* was the effective percentage of the distribution frequency within the material at level *i* of a trait[15].

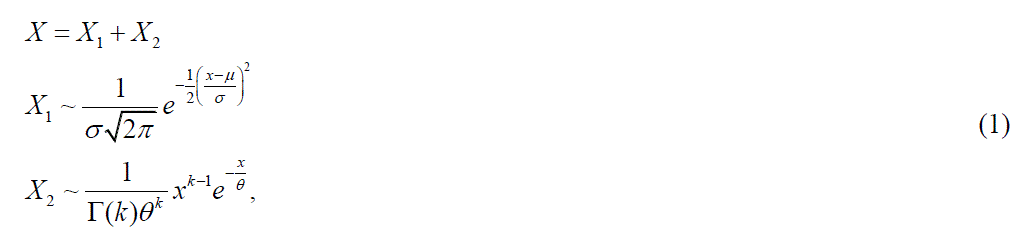

V_ST_ was the phenotypic differentiation coefficient, representing the percentage of inter population variation of total genetic variation.

δ_2t/S_ was the inter-group variance component, and δ_2S_ was the intra-group variance component [16].

## Results

### Genetic diversity analysis of fruit phenotypic traits of ancient *Cinnamomum camphora* in different provenances

The differences of fruit phenotypic traits were showed in Table 2. The differences among provenances of FTW, FV, FHD, PT, FSD/FHD, FSD/FVD reached a extremely significant level (*P*<0.01). There were significant differences of FTW, FSD/FHD, FSD/FVD within provenances (*P*<0.05). The variance component among provenances(20.35%) was greater than within provenances(5.306%), indicating that variation within provenances was the main source in ancient *Cinnamomum camphora* in Jiangxi Province, China. The variation range of phenotypic differentiation coefficient among provenances was 65.6% to 100%, all more than 50%. The phenotypic differentiation coefficient of FTW/FV was the highest, while phenotypic differentiation coefficient of FVD, FV and FSD were all more than 80%.

**Table 2.** Comparison of phenotypic traits and phenotypic differentiation coefficients of ancient *Cinnamomum camphora* fruits from different provenances. * and ** indicate significant difference at *P*<0.05 and *P*<0.01, respectively. FTW, Fruit thousand-grain weight; FV, Fruit volume; FSD, Fruit side diameter; FHD, Fruit horizontal diameter; FVD, Fruit vertical diameter; PT, Peel thickness.

| Fruit phenotypic character | Among provenances |  |  |  | Within provenances |  |  |  | Random error |  |
| --- | --- | --- | --- | --- | --- | --- | --- | --- | --- | --- |
|  | Mean square | F value | Heft /% | Phenotypic differentiation coefficient/% | Mean square | F value | Heft /% | Phenotypic differentiation coefficient/% | Mean square | Heft /% |
| **FTW** | 30683.952 | 4.112** | 16.81 | 65.6 | 8094.2 | 1.085* | 8.78 | 34.31 | 7462.915 | 74.41 |
| **FV** | 29588.821 | 4.211** | 17.98 | 83.2 | 7550.89 | 1.075 | 3.64 | 16.83 | 7027.192 | 78.37 |
| **FSD** | 0.656 | 1.716 | 17.72 | 84.0 | 0.322 | 0.844 | 3.36 | 15.9 | 0.382 | 78.92 |
| **FHD** | 0.704 | 2.303** | 16.14 | 72.5 | 0.28 | 0.915 | 6.1 | 27.4 | 0.306 | 77.76 |
| **FVD** | 0.648 | 1.841 | 27.51 | 94.4 | 0.253 | 0.717 | 1.61 | 5.5 | 0.352 | 70.8 |
| **PT** | 0.259 | 3.755** | 34.84 | 80.8 | 0.085 | 1.225 | 8.27 | 19.2 | 0.069 | 56.89 |
| **FSD/FHD** | 0.002 | 3.276** | 23.24 | 77.8 | 0.001 | 1.341* | 6.61 | 22.2 | 0.001 | 70.15 |
| **FSD/FVD** | 0.008 | 3.248** | 17.94 | 66.3 | 0.003 | 1.261* | 9.1 | 33.7 | 0.002 | 72.96 |
| **FHD/FVD** | 0.005 | 2.615 | 17.85 | 76.2 | 0.002 | 0.998 | 5.59 | 23.8 | 0.002 | 76.56 |
| **FTW/FV** | 0.001 | 0.65 | 13.44 | 100 | 0.002 | 1.193 | 0 | 0 | 0.001 | 86.56 |
| **Average** | 6027.506 | 2.773 | 20.35 | 80.1 | 1564.60 | 1.0654 | 5.31 | 19.89 | 1449.122 | 74.34 |

\* and \*\* indicate significant difference at $P<0.05$ and $P<0.01$ , respectively.

There were significant differences of fruit phenotypes of ancient *Cinnamomum camphora* from 11 provenances in Jiangxi, China (Table 3), and the variation ranges of FTW, FV and PT were 454.9-608.72g, 438.19-586.43 cm^3^, and 0.89-1.28 mm,respectively. The FHD, FVD and FSD were 8.09-9.08 mm, 8.46-9.36 mm, and 8.91-9.77 mm. The FSD/FHD, FSD/FVD, FHD/FVD and FTW/FV were 1.05 to 1.11, 1.02 to 1.47, 0.94 to 1.35, and 1.03 to 1.06g·cm^-3^, respectively. The Shannon Wiener index of 10 traits were ranged from 2.40 to 2.85, among which FHD/FVD was the highest and FTW, FV were the lowest. It was indicated that the fruits of ancient *Cinnamomum camphora* from different provenances had rich genetic diversity.

**Table 3.** Comparison on fruit phenotypic characters of ancient *Cinnamomum camphora* from different provenances and analysis on hannon-Wiener index (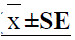)

| prov<br>enan<br>ce | FTW/g | FV/cm<sup>3</sup> | FSD/mm | FHD/mm | FVD/mm | PT/mm | FSD<br>/FHD | FSD<br>/FVD | FHD<br>/FVD | FTW/FV<br>(g·cm<sup>-3</sup>) |
| --- | --- | --- | --- | --- | --- | --- | --- | --- | --- | --- |
| NC | 488.65±33.72bc | 468.73±34.86bcd | 9.07±0.35bc | 8.2±0.31bc | 8.76±0.32ab | 0.98±0.11bc | 1.11±0.01a | 1.03±0.00ab | 0.94±0.01ab | 1.05±0.01a |
| YT | 478.52±33.26bc | 451.85±30.32cd | 8.91±0.22c | 8.09±0.23c | 8.46±0.24b | 1.02±0.04abc | 1.1±0.01a | 1.05±0.02ab | 0.96±0.02ab | 1.06±0.00a |
| SR | 514.2±12.56bc | 490.85±12.06bcd | 9.33±0.11abc | 8.63±0.09abc | 9.15±0.09a | 1.27±0.04ab | 1.08±0.00ab | 1.02±0.01ab | 0.94±0.01ab | 1.05±0.00a |
| FZ | 552.79±23.61ab | 534.81±20.94abc | 9.26±0.15abc | 8.49±0.13abc | 8.78±0.18ab | 0.99±0.06bc | 1.09±0.01ab | 1.47±0.41a | 1.35±0.38a | 1.03±0.01a |
| JJ | 454.94±17.22c | 438.19±14.08d | 9.2±0.09abc | 8.61±0.08abc | 8.99±0.07ab | 0.89±0.09c | 1.07±0.00b | 1.02±0.01ab | 0.96±0.01ab | 1.04±0.01a |
| YC | 570.01±19.24ab | 546.71±19.44ab | 9.66±0.14ab | 8.81±0.13a | 8.91±0.13ab | 1.13±0.05abc | 1.1±0.01ab | 1.06±0.02ab | 0.99±0.01ab | 1.03±0.01a |
| JDZ | 474.57±22.10bc | 460.32±21.53bcd | 9.54±0.22abc | 8.75±0.13ab | 9.1±0.25a | 1.28±0.04a | 1.09±0.01ab | 1.05±0.03ab | 0.96±0.02ab | 1.03±0.02a |
| JA | 608.72±16.47a | 586.43±15.91a | 9.77±0.11a | 9.08±0.09a | 9.36±0.13a | 1.28±0.07a | 1.08±0.00ab | 1.05±0.01ab | 0.97±0.01ab | 1.04±0.00a |
| PX | 548.22±39.93abc | 526.36±39.51abcd | 9.62±0.21ab | 8.87±0.20a | 9.27±0.17a | 1.1±0.08abc | 1.09±0.01ab | 1.04±0.02ab | 0.96±0.01ab | 1.04±0.01a |
| XY | 539.75±31abc | 522.22±34.38abcd | 9.27±0.23abc | 8.86±0.13a | 8.93±0.13ab | 1.28±0.14a | 1.05±0.01c | 1.04±0.02ab | 0.99±0.01ab | 1.04±0.01a |
| GZ | 565.05±16.91ab | 544.09±16.64abc | 9.55±0.11abc | 8.81±0.11a | 9.25±0.10a | 1.13±0.05abc | 1.08±0.01ab | 1.03±0.01ab | 0.95±0.01ab | 1.04±0.01a |
| H' | 2.395 | 2.395 | 2.446 | 2.45 | 2.448 | 2.795 | 2.817 | 2.813 | 2.848 | 2.835 |

Different lower cases in the same column indicate the significant difference at 0.05 level.
H' stands for Shannon-Wiener index.

Variation coefficient of phenotypic traits in different provenances was shown in table 4. Among 10 fruit phenotypic traits in different provenances, the largest coefficient of variation was PT(25.45%) followed by FV as the second highest (17.83%), and the smallest coefficient of variation was FSD/FHD(2.64%). According to the mean value of coefficient variation of phenotypic traits in 11 provenances, 2 provenances were more than 10%, among which the Pingxiang provenance was the largest with the value for 11.14%, followed by Nanchang provenance (10.68%), and the Yingtan provenance was the smallest(5.15%). The mean value of the coefficient of variation among provenances (9.65%) was greater than the variation within provenances (8.52%) for fruit phenotypic traits of ancient *Cinnamomum camphora*.

**Table 4.** Analysis of coefficient variation of fruit phenotypic traits of ancient *Cinnamomum camphora* from different provenances in Jiangxi Province.

| Provenances | FTW | FV | FSD | FHD | FVD | PT | FSD /FHD | FSD /FVD | FHD /FVD | FTW /FV | Average |
| --- | --- | --- | --- | --- | --- | --- | --- | --- | --- | --- | --- |
| NC | 18.26 | 19.68 | 10.14 | 10.01 | 9.65 | 28.32 | 3.13 | 1.23 | 2.96 | 3.39 | 10.68 |
| YT | 12.04 | 11.62 | 4.23 | 4.90 | 4.94 | 6.59 | 0.84 | 2.51 | 3.11 | 0.69 | 5.15 |
| SR | 13.38 | 13.46 | 6.53 | 5.93 | 5.65 | 16.55 | 2.48 | 4.08 | 3.33 | 2.12 | 7.35 |
| FZ | 16.54 | 15.16 | 6.10 | 6.00 | 8.03 | 22.19 | 2.88 | 6.43 | 5.89 | 3.06 | 9.23 |
| JJ | 15.14 | 12.85 | 3.92 | 3.58 | 3.32 | 41.57 | 1.71 | 4.04 | 3.80 | 4.30 | 9.42 |
| YC | 17.54 | 18.47 | 7.27 | 7.65 | 7.38 | 25.20 | 2.46 | 4.32 | 4.08 | 3.99 | 9.84 |
| JDZ | 12.32 | 12.38 | 6.03 | 4.05 | 7.20 | 8.61 | 3.18 | 7.70 | 6.75 | 5.91 | 7.41 |
| JA | 11.79 | 11.82 | 4.71 | 4.52 | 6.00 | 23.11 | 1.65 | 5.24 | 4.38 | 1.90 | 7.51 |
| PX | 24.15 | 24.89 | 7.39 | 7.67 | 6.12 | 25.96 | 3.12 | 5.08 | 5.08 | 1.97 | 11.14 |
| GZ | 16.66 | 17.03 | 6.64 | 6.67 | 6.04 | 24.59 | 2.70 | 4.74 | 4.50 | 4.65 | 9.42 |
| CV/% | 17.66 | 17.83 | 6.64 | 6.56 | 6.57 | 25.45 | 2.64 | 5.08 | 4.56 | 3.51 | / |

### Correlation analysis on fruit phenotypic traits of ancient *Cinnamomum camphora* in Jiangxi Province

There were certain correlations among fruit phenotypic traits of ancient *Cinnamomum camphora* from different provenances in Jiangxi Province, China(Fig 1). There were significant positive correlations between FTW and four traits of FV, FSD, FHD and FVD(*P*<0.05). FV showed a highly significant correlation with FSD, FHD and FVD.

**Fig 1.**
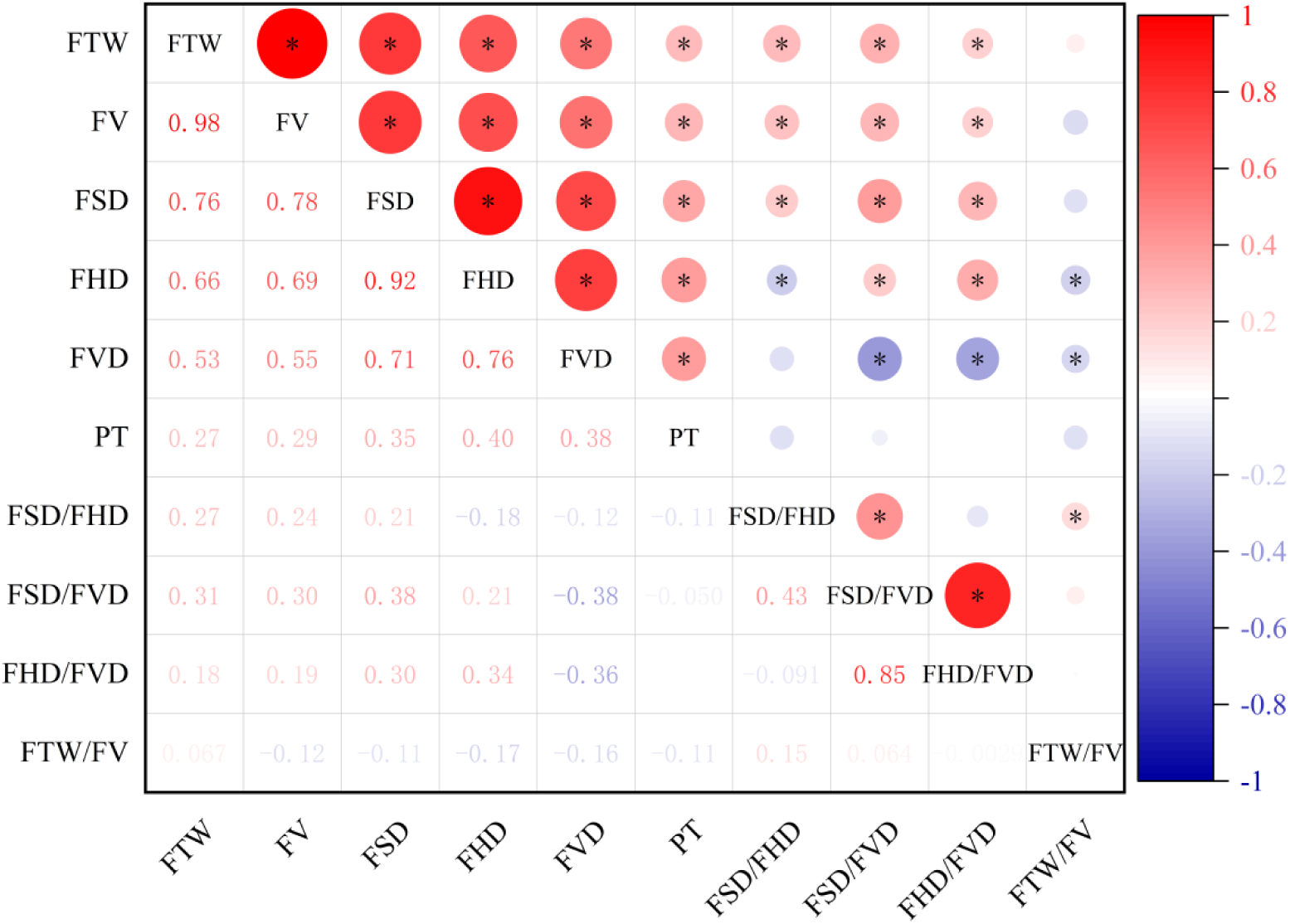
Correlation analysis among 10 fruit phenotypic traits of ancient *Cinnamomum camphora*.

FSD showed significantly positively correlated with FHD and FV. FHD was significantly positively correlated with FV while negatively correlated with FSD/FHD and FTW/FV. FVD showed significantly positively correlated with PT while there were significant negative correlations with FSD/FVD, FHD/FVD, and FTW/FV. In summary, most of the relevant traits of ancient *Cinnamomum camphora* fruits were closely related, and when one trait was changed, it might cause changes in other indexes.

### Response of fruit phenotypic traits to longitude and climatic factors from different provenances in Jiangxi Province

The correlation analysis between the phenotypic traits of ancient *Cinnamomum camphora* from different provenances in Jiangxi Province and the latitude and longitude were shown in Figs 2 and 3. FTW and FV showed a highly significant negative correlation with latitude(*P*<0.01), while FSD and FHD showed a significant negative correlation with latitude (*P*<0.05). FTW and FV showed a highly significant negative correlation with latitude. FSD, FHD, FSD/FVD and FHD/FVD showed a significant negative correlation with longitude, which indicated that there were significant geographic variations in the phenotypic traits of ancient *Cinnamomum camphora* in Jiangxi Province.

**Fig 2.**
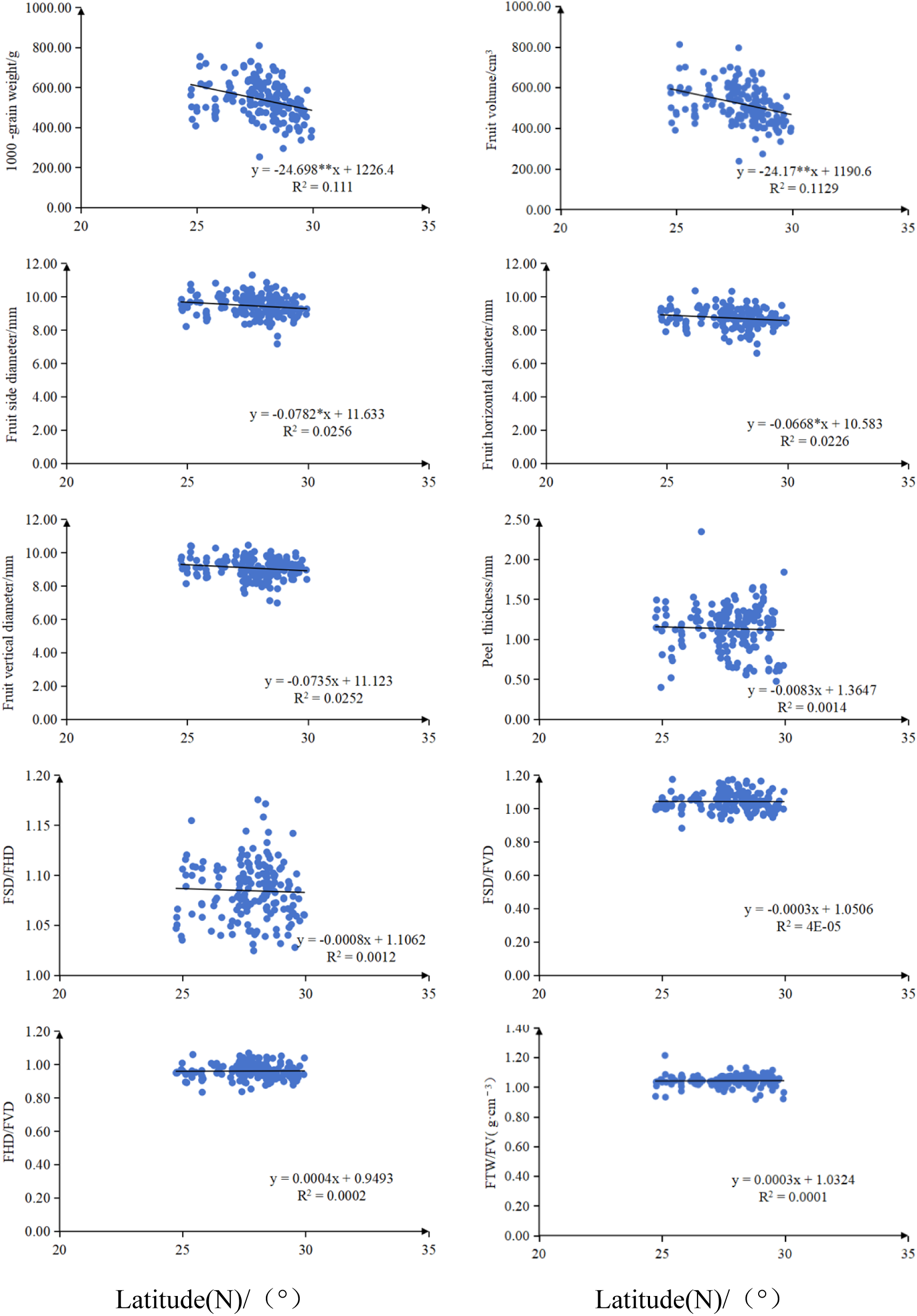
Relationship between fruit traits of ancient *Cinnamomum camphora* from different provenances in Jiangxi Province and latitude.

**Fig 3.**
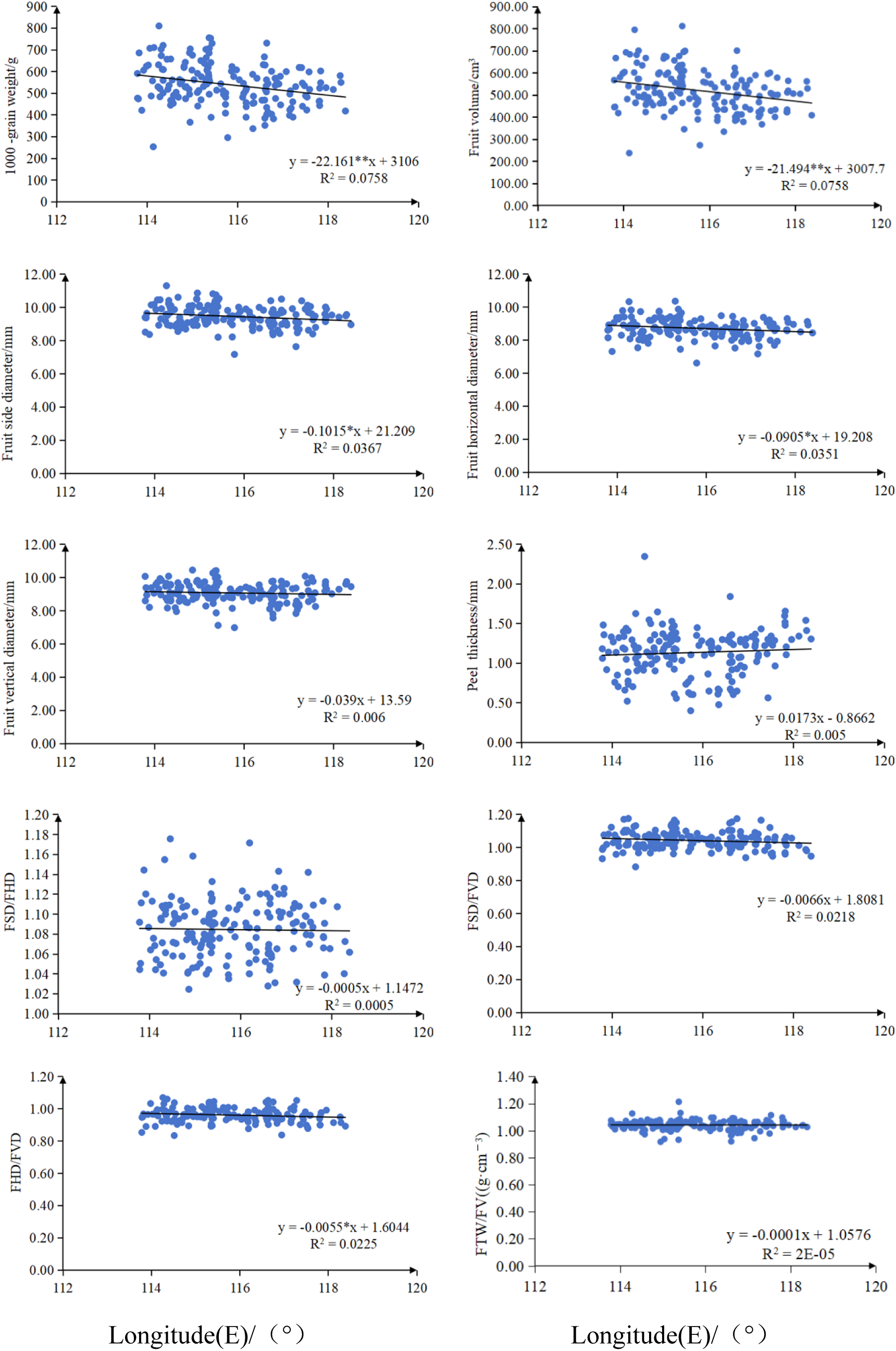
Relationship between fruit traits of ancient *Cinnamomum camphora* from different provenances in Jiangxi Province and longitude.

The results of the correlation analysis between fruits phenotypic traits of ancient *Cinnamomum camphora* and Geo-climatic factors in Jiangxi Province were shown in Table 5. FTW and FV were positively correlated with the mean temperature in January, mean altitude(*P*<0.01) and annual mean temperature(*P*<0.05), while negatively correlated with annual average sunshine duration(*P*<0.01) and mean temperature in July(*P*<0.05). FSD was negatively correlated with annual average sunshine duration(*P*<0.01). And FHD was significantly negatively correlated with mean temperature in July and annual average sunshine duration(*P*<0.05). FVD was positively correlated with mean altitude(*P*<0.05). PT was positively correlated with mean altitude and annual mean precipitation(*P*<0.05). FSD/FHD was positively correlated with annual mean precipitation(*P*<0.05). FSD/FVD was negatively correlated with annual average sunshine duration(*P*<0.05).

**Table 5.** Correlation between fruit phenotypic traits of ancient *Cinnamomum camphora* and climatic factors.

| Fruit phenotypic trait | Annual mean temperature /°C | Mean temperature in January/°C | Mean temperature in July/°C | Annual average sunshine duration/h | Mean altitude/m | Annual mean precipitation /mm |
| --- | --- | --- | --- | --- | --- | --- |
| **FTW** | 1.77* | 0.216** | -0.162* | -0.374** | 0.297** | 0.006 |
| **FV** | 0.178* | 0.218** | -0.169* | -0.371** | 0.290** | -0.003 |
| **FSD** | 0.037 | 0.081 | -0.150 | -0.205** | 0.134 | -0.039 |
| **FHD** | 0.015 | 0.078 | -0.164* | -0.174* | 0.119 | -0.112 |
| **FVD** | 0.109 | 0.137 | -0.143 | 0.109 | 0.166* | -0.103 |
| **PT** | 0.120 | 0.100 | 0.088 | -0.079 | 0.187* | 0.195* |
| **FSD/FHD** | 0.066 | 0.016 | 0.029 | -0.079 | 0.051 | 0.187* |
| **FSD/FVD** | -0.088 | -0.064 | -0.019 | -0.173* | -0.035 | 0.088 |
| **FHD/FVD** | -0.135 | -0.083 | -0.030 | -0.137 | -0.069 | 0.002 |
| **FTW/FV** | -0.009 | -0.016 | 0.051 | -0.018 | 0.037 | 0.057 |

### Principal component analysis of fruit phenotypic traits of different provenances in Jiangxi Province

Principal component analysis of fruit phenotypic traits from different provenances in Jiangxi Province of China, in which three principal components with eigenvalues greater than 1.00 were yielded with a cumulative contribution rate of 78.78%. It was indicated that the three principal components represented most of the information on phenotypic traits of ancient *Cinnamomum camphora* (Table 6). Among them, the 1st principal component had the highest contribution rate of 42.62%, with an eigenvalue for 4.26, and eigenvector absolute value of FSD (0.96), FHD (0.91), FV (0.91) and FTW (0.89) were larger, which indicated that the 1st principal component was mainly associated with the traits related to fruit size. The contribution rate of the 2nd principal component was 22.55% with the eigenvalue for 2.255, and the absolute values of 3 trait eigenvectors of FSD/FVD (0.92), FHD/FVD (0.78), and FVD (–0.67) were larger, indicating that the 2nd principal component was related to fruit shape. The contribution rate of the 3rd principal component was 13.61%, and the eigenvalue was 1.361, in which FSD/FHD (0.76) and FHD/FVD (–0.54) had the largest absolute value, indicating that the 3rd principal component was also a trait mainly reflecting fruit shape. In summary, FSD, FSD/FVD and FSD/FHD could comprehensively reflect the information of 10 traits.

**Table 6.** The principal component analysis of ancient *Cinnamomum camphora* fruit traits in different provenances of Jiangxi Province.

| Character index | First principal component | Second principal component | Third principal component |
| --- | --- | --- | --- |
| **FSD** | 0.957 | 0.034 | 0.006 |
| **FHD** | 0.907 | -0.137 | -0.292 |
| **FV** | 0.906 | 0.043 | 0.200 |
| **FTW** | 0.885 | 0.086 | 0.293 |
| **FVD** | 0.694 | -0.671 | 0.091 |
| **PT** | 0.442 | -0.304 | -0.225 |
| **FSD/FVD** | 0.344 | 0.921 | -0.100 |
| **FHD/FVD** | 0.294 | 0.775 | -0.540 |
| **FSD/FHD** | 0.143 | 0.429 | 0.760 |
| **FTW/FV** | -0.132 | 0.224 | 0.460 |
| **Eigenvalue** | 4.262 | 2.255 | 1.361 |
| **Contribution rate%** | 42.617 | 22.553 | 13.609 |
| **Cumulative contribution rate%** | 42.617 | 65.170 | 78.779 |

According to the component factor score coefficient matrix in Table 6, three principal component factor scores could be calculated by using the standardized quality indexes. The calculation formula of factor score of each component was as follows: y_1_=0.464x_1_+0.439x_2_+0.439x_3_+0.429x_4_+0.336x_5_+0.214x_6_+0.167x_7_+0.142x_8_+0.069x_9_-0.064x_10_ y_2_=0.023x_1_-0.091x_2_+0.029x_3_+0.057x_4_-0.447x_5_-0.202x_6_+0.613x_7_+0.516x_8_+0.286x_9_+0.149x_10_ y_3_=0.005x_1_-0.250x_2_+0.171x_3_+0.251x_4_+0.078x_5_-0.193x_6_-0.086x_7_-0.463x_8_+0.651x_9_+0.394x_10_ Where x_1_ and x_10_ represented the standardized indicators of 10 fruit phenotypic traits, y_1_, y_2_ and y_3_ were the main components 1st, 2nd and 3rd respectively. The characteristic values of principal components 1st, 2nd and 3rd were divided by the sum of characteristic values. According to comprehensive quality score function, the formula was as follows.

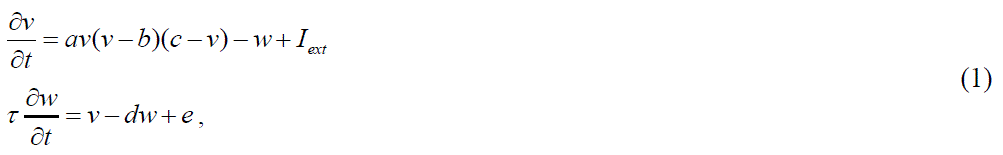

According to fruit trait load coefficient of ancient *Cinnamomum camphora* on the three principal components and the contribution rate of the three principal components, the comprehensive score of each ancient *Cinnamomum camphora* tree was calculated, so as to obtain the top 20 plants of ancient *Cinnamomum camphora* in the comprehensive score, with the sample number for 79, 67, 112, 166, 72, 137, 98, 165, 142, 120, 66, 159, 80, 97, 87, 11, 59, 135, 109 and 157 and the corresponding scores were 3.08, 2.12, 1.82, 1.81, 1.77, 1.76, 1.60, 1.59, 1.56, 1.53, 1.52, 1.483, 1.47, 1.29, 1.25, 1.24, 1.19, 1.18, 1.15 and 1.12. Among them, there were 6 strains from Yichun, 5 from Ganzhou, 4 from Ji’an, 2 from Fuzhou, 2 from Pingxiang, and 1 from Jingdezhen (Table 7).

**Table 7.**
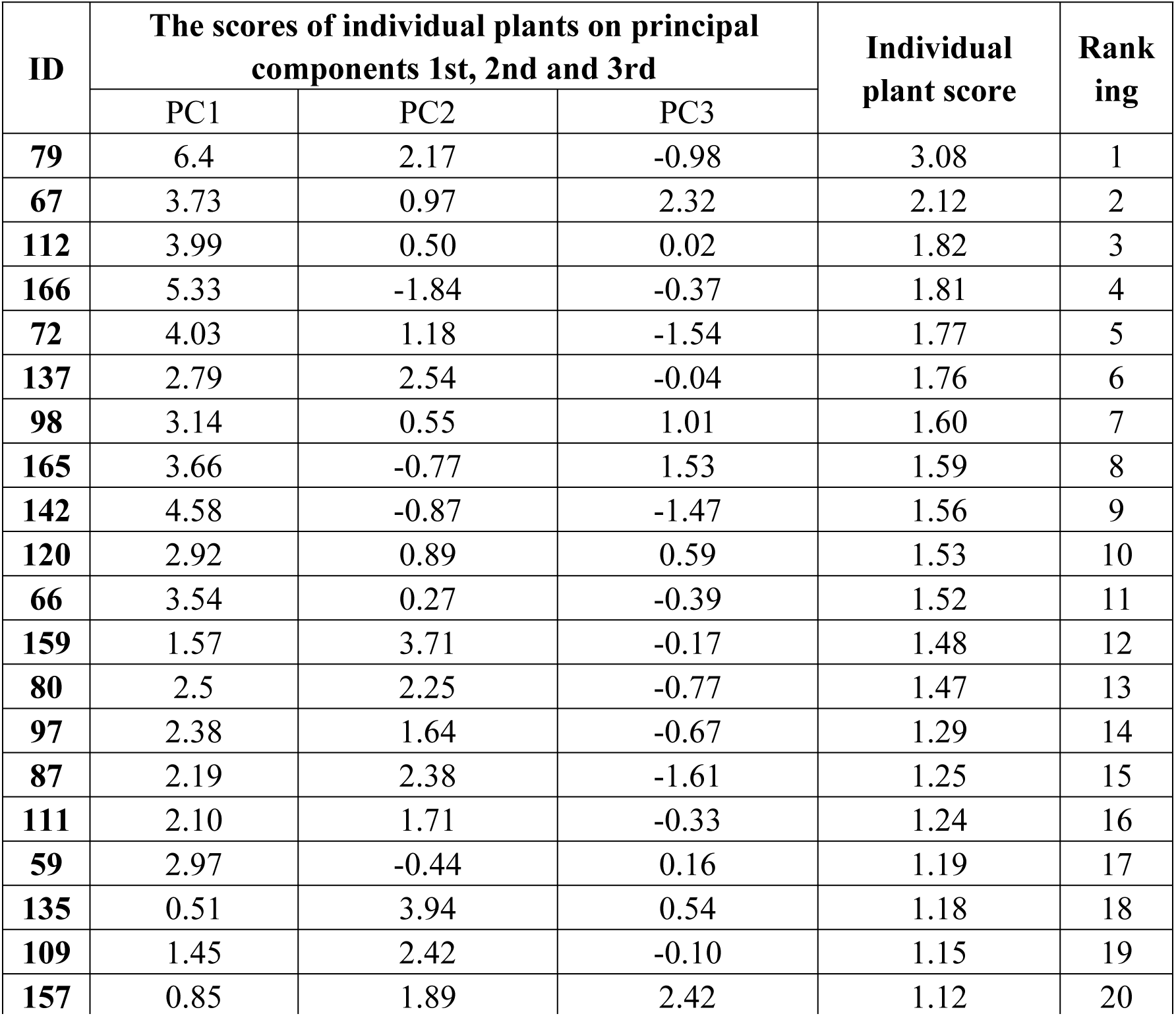
Principal component scores and comprehensive scores of the top 20 individual ancient *Cinnamomum camphora* plants with comprehensive scores.

| ID | The scores of individual plants on principal components 1st, 2nd and 3rd |  |  | Individual plant score | Ranking |
| --- | --- | --- | --- | --- | --- |
|  | PC1 | PC2 | PC3 |  |  |
| 79 | 6.4 | 2.17 | -0.98 | 3.08 | 1 |
| 67 | 3.73 | 0.97 | 2.32 | 2.12 | 2 |
| 112 | 3.99 | 0.50 | 0.02 | 1.82 | 3 |
| 166 | 5.33 | -1.84 | -0.37 | 1.81 | 4 |
| 72 | 4.03 | 1.18 | -1.54 | 1.77 | 5 |
| 137 | 2.79 | 2.54 | -0.04 | 1.76 | 6 |
| 98 | 3.14 | 0.55 | 1.01 | 1.60 | 7 |
| 165 | 3.66 | -0.77 | 1.53 | 1.59 | 8 |
| 142 | 4.58 | -0.87 | -1.47 | 1.56 | 9 |
| 120 | 2.92 | 0.89 | 0.59 | 1.53 | 10 |
| 66 | 3.54 | 0.27 | -0.39 | 1.52 | 11 |
| 159 | 1.57 | 3.71 | -0.17 | 1.48 | 12 |
| 80 | 2.5 | 2.25 | -0.77 | 1.47 | 13 |
| 97 | 2.38 | 1.64 | -0.67 | 1.29 | 14 |
| 87 | 2.19 | 2.38 | -1.61 | 1.25 | 15 |
| 111 | 2.10 | 1.71 | -0.33 | 1.24 | 16 |
| 59 | 2.97 | -0.44 | 0.16 | 1.19 | 17 |
| 135 | 0.51 | 3.94 | 0.54 | 1.18 | 18 |
| 109 | 1.45 | 2.42 | -0.10 | 1.15 | 19 |
| 157 | 0.85 | 1.89 | 2.42 | 1.12 | 20 |

### Cluster analysis based on fruit phenotypic traits from different provenances of ancient *Cinnamomum camphora* in Jiangxi Province

Distance cluster analysis was used to analyze the 10 fruit phenotypic traits from 11 provenances of ancient *Cinnamomum camphora* in Jiangxi Province of China (Fig 4), and the mean values of fruit traits for each group were calculated (Table 8). The results showed that ancient *Cinnamomum camphora* from different provenances could be divided into 2 group at the distance coefficient of 1.2. At the same time, they were further divided into different sub groups at the distance coefficient of 0.9, of which A was a separate group and B was divided into B1 and B2.

**Figure 4.**
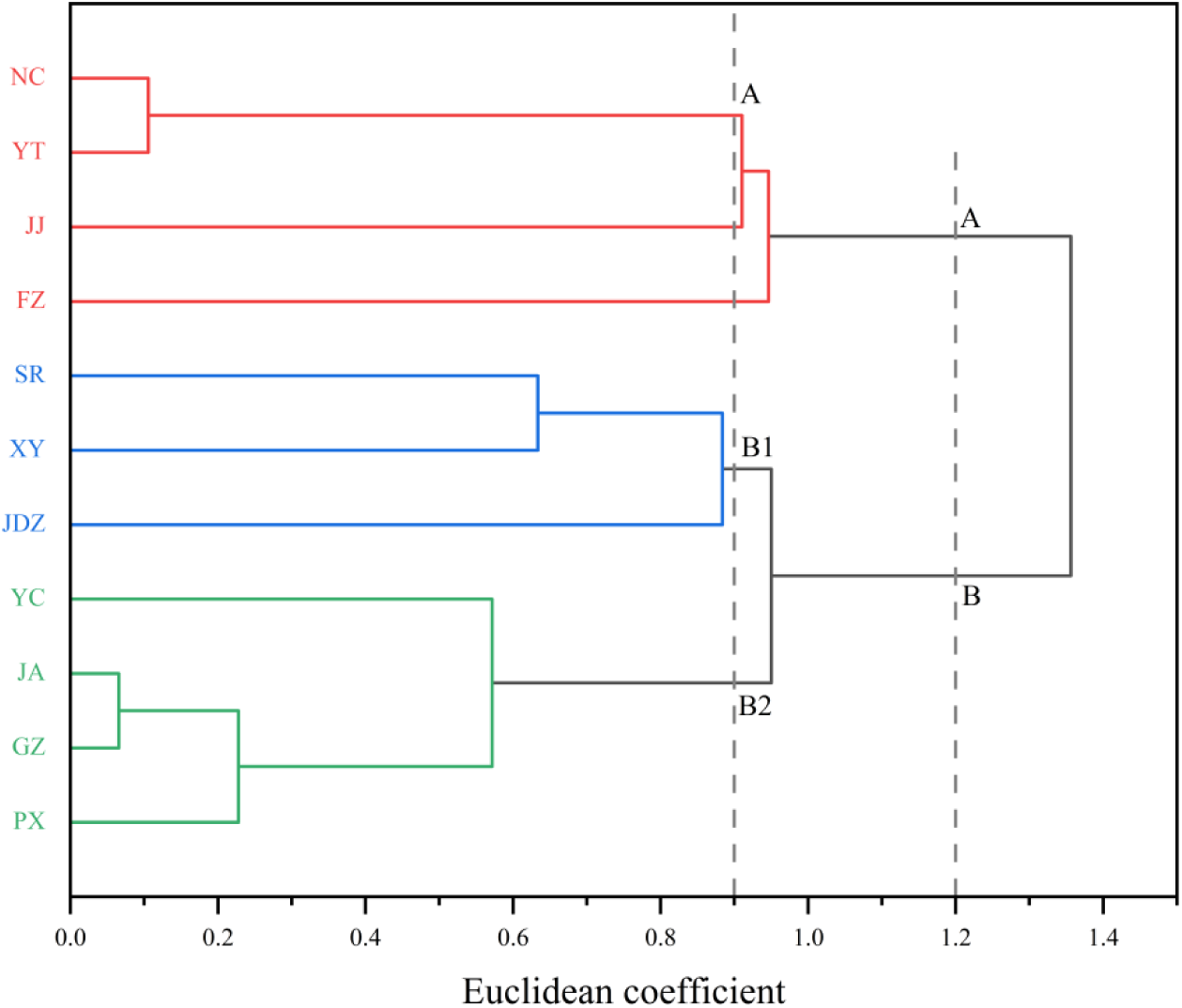
The cluster analysis of fruit traits of ancient *Cinnamomum camphora* from different provenances.

**Table 8.** Comparison of fruit phenotypic traits of different groups of ancient *Cinnamomum camphora*.

| Group | FTW/g | FV/cm<sup>3</sup> | FSD/mm | FHD/mm | FVD/mm | PT/mm | FSD/FHD | FSD/FVD | FHD/FVD | FTW/FV (g·cm<sup>-3</sup>) |
| --- | --- | --- | --- | --- | --- | --- | --- | --- | --- | --- |
| **Group** |  |  |  |  |  |  |  |  |  |  |
| **A** | 493.72b | 473.40b | 9.11b | 8.35b | 8.75b | 0.97b | 1.09a | 1.14a | 1.05a | 1.04a |
| **B** | 545.79a | 525.28a | 9.53a | 8.83a | 9.14a | 1.21a | 1.08b | 1.04b | 0.97b | 1.04a |
| **Subgroup** |  |  |  |  |  |  |  |  |  |  |
| **A** | 493.725b | 473.395b | 9.11c | 8.348b | 8.75b | 9.7c | 1.093a | 1.143a | 1.053a | 1.05a |
| **B1** | 509.507b | 491.130b | 9.38b | 8.747a | 9.06ab | 1.28a | 1.037a | 1.037a | 0.963a | 1.04a |
| **B2** | 573.00a | 550.898a | 9.65a | 8.893a | 9.196a | 1.16b | 1.045a | 1.045a | 0.968a | 1.038a |

Group A included four provenances from Nanchang, Yingtan, Jiujiang and Fuzhou. Subgroup B1 included the three provenances of Shangrao, Jingdezhen and Xinyu, and Subgroup B2 was the four provenances of Ji’an, Yichun, Ganzhou and Pingxiang. The fruit phenotypic traits of group B provenances were significantly greater than those of group A, as evidenced by thicker peel, larger fruits, and heavier fruit. FTW, FV and FSD in the B2 provenances were significantly higher than B1, while PT in the B2 provenances was significantly lower than B1 subgroup.

## Discussion

### Diversity and variation of fruit phenotypic traits of ancient *Cinnamomum camphora*

Ancient *Cinnamomum camphora* have a wide natural distribution area. Complex habitats and geographic isolation have resulted in genetic differentiation of phenotypic traits among different provenances. Phenotypic variation is the most direct manifestation of biological genetic variation, and it is also one of the important contents of genetic diversity research[17–18]. Carrying out studies on diversity of fruit phenotypic traits, to some extent, can reflect the variation and the level of genetic diversity in different populations [19–20].

The present study is shown that there is abundant variation in fruit phenotypic traits of ancient *Cinnamomum camphora* in Jiangxi Province, both among and within the provenances, and the variation mainly originates from among provenances, which is the result of the genetic factors of *Cinnamomum camphora* themselves and the long-term environmental heterogeneity. The average value of the phenotypic differentiation coefficient of ancient *Cinnamomum camphora* fruits among different provenances was 80.08%, which was significantly higher than that of *Xanthoceras sorbifolium* Bunge, *Quercus fabri* Hance and *Armeniaca mandshurica* (Maxim.) Skv.[21–23], indicating a high degree of phenotypic differentiation of ancient *Cinnamomum camphora* from different provenances. Meanwhile, the Shannon-Wiener index of fruit phenotypic traits of ancient *Cinnamomum camphora* in Jiangxi Province was fluctuating in the range of 2.395 to 2.835, which was close to that of *Zanthoxylum armatum* DC.[24]. However, the phenotypic diversity indices of different plants were different, such as *Capsicum annuum* L.varied in the range of 5. 177 to 5.368[25], whereas the values of *Ziziphus jujuba* Mill.(1.04 to 2.05) and *Juglans cathayensis* Dode(1.41 to 2.02) were smaller[26–27]. Ancient *Cinnamomum camphora* was predominantly heterogamous, and artificial planting, introductions would increase the degree of variation, which can affect the differences in fruit phenotypes. In this study, ancient *Cinnamomum camphora* were selected as the research object to reduce the variability caused by artificial introduction.

The coefficient of variation is an index reflecting the degree of plant variation. It is generally recognized that coefficient variation higher than 10% indicates a large variation in plant traits [28–29]. In this study, the average coefficient variation of fruit phenotypic traits in ancient *Cinnamomum camphora* among different provenances in Jiangxi Province was 9.65%, of which the coefficient of variation of PT was the largest, which was consistent with the results of Li et al.[30] and Gao et al.[31]. The average value of coefficient variation of fruit phenotypes within provenances in Jiangxi Province was 5.15%-11.14%, which was consistent with the result of Hunan Province[32]. The average coefficient variation of FTW and FV of ancient *Cinnamomum camphora* in this study (16.11%) was much larger than that of morphological indicators such as FVD and FHD (5.34%), which indicated that fruit weights and volumes in ancient *Cinnamomum camphora* were more stable, which was in agreement with the results of Xing et al.[33].

### Geographical variation and clustering results of fruit phenotypic traits in ancient *Cinnamomum camphora*

The geographic pattern of variation in plant fruit phenotypic traits is complex and is usually determined by a combination of genotypes and environmental factors. Different plants have different patterns of variation, such as *Rhus chinensis* Mill. showed a pattern of variation dominated by frost-free period and mean annual temperature[12], while *Armeniaca vulgaris* var. ansu showed a pattern of variation dominated by altitude[34].The correlation between fruit phenotypic traits of *Cinnamomum camphora* from Jiangxi Province and geographic and climatic factors in this study was strong, and most of the fruit phenotypic traits were significantly or highly significantly positively correlated with annual mean temperature, mean temperature in January, and mean altitude. Previous studies have shown that plant fruit size decreases with increasing altitude, possibly due to environmental changes that cause a plastic response of plant to the reduction of available resources[35]. However, the results of the present study were contrary to this, with increasing altitude, FTW, FV, and PT were greater. As the altitude range of the sampling sites was 32-388 m, the increase in altitude could alleviate the stress of high temperature to a certain extent and increase the temperature difference between day and night, which was favorable to fruit expansion and nutrient accumulation. In some regions, plant fruits become heavier with increasing sunlight, such as Yuan et al.[36], who studied the fruit trait characteristics of *Magnolia wilsonii* (Finet et Gagn) Rehd. and its phenotypic diversity and found that the mass of individual polymerized follicles was highly significantly and positively correlated with the annual average sunlight, but in the present study the fruit phenotypes in ancient *Cinnamomum camphora* were significantly or highly significantly negatively correlated with the annual average sunshine duration and the mean temperature in July. Jiangxi is located in subtropical monsoon climate, high temperature and rainy in summer. The increase of sunshine hours will exacerbate the high temperature and heat wave stress to a certain extent, and the fruit growth and development is poor under high temperature stress. In addition, FTW, FV and FSD, FHD and other morphological indicators of ancient *Cinnamomum camphora* from different provenances in Jiangxi Province had a decreasing trend with the rise of latitude and longitude, indicating that the phenotypic traits of ancient *Cinnamomum camphora* were affected by the effects of changes in temperature and precipitation caused by changes in geographic location gradient.

Principal component analysis can comprehensively analyze multiple traits in order to obtain more intuitive and simpler results, and at the same time exclude the interference of complex interrelationships among multiple trait indicators on the analysis of results[37]. In this study, it was found that three indicators of FSD, FSD/FVD and FSD/FHD could comprehensively reflect the 10 phenotypic traits of ancient *Cinnamomum camphora* in Jiangxi Province, which could be used as an important basis for judging the growth status of the ancient *Cinnamomum camphora*. In addition, 20 single plants of ancient *Cinnamomum camphora* screened from different provenances in Jiangxi Province had larger and heavier fruits, which could provide certain reference for the selection of germplasm resources of ancient *Cinnamomum camphora* in Jiangxi Province.

According to the results of clustering analysis of fruit phenotypic traits of ancient *Cinnamomum camphora* from different provenances in this study, it was found that the 11 provenances in Jiangxi Province could be divided into three groups, of which the fruits from B2 group were heavier and larger in fruit morphology, which were more accordance with the standard of artificially selecting excellent germplasm resources. However, the provenances were not completely clustered according to the geographical distance, probably because of the influence of the differences in soil, topography and other comprehensive fact.

## Conclusions

The fruit phenotypic traits of ancient *Cinnamomum camphora* from different provenances in Jiangxi Province contained rich variations, and the variations were mainly originated from among provenances. FTW, FV, and FSD all differed significantly among provenances. FSD, FSD/FVD and FSD/FHD could be used as the main indexes to comprehensively reflect fruit phenotypic traits of ancient *Cinnamomum camphora.* There was a significant correlation between fruit phenotypic traits and latitude, longitude, climate factors. According to the results of cluster analysis, ancient *Cinnamomum camphora* from 11 provenances could be divided into three major groups, in which the fruit phenotypic traits of B2 provenances were larger. It will provide an important direction for the selection of excellent germplasm resources of ancient *Cinnamomum camphora*.

## Notes

### Competing Interest Statement

The authors have declared no competing interest.

